# An intranasal stringent response vaccine targeting dendritic cells as a novel adjunctive therapy against tuberculosis

**DOI:** 10.1101/2022.04.19.488816

**Authors:** Styliani Karanika, James T. Gordy, Pranita Neupane, Theodoros Karantanos, Jennie Ruelas Castillo, Darla Quijada, Kaitlyn Comstock, Avinaash Kaur Sandhu, Yinan Hui, Samuel K. Ayeh, Rokeya Tasneen, Stefanie Krug, Carina Danchik, Tianyin Wang, Courtney Schill, Rirchard B. Markham, Petros C. Karakousis

## Abstract

Lengthy tuberculosis (TB) treatment is required to address the ability of a subpopulation of persistent *Mycobacterium tuberculosis* (*Mtb*) to remain in a non-replicating, antibiotic-tolerant state characterized by metabolic remodeling, including induction of the Rel_Mtb_-mediated stringent response. We developed a novel therapeutic DNA vaccine construct involving fusion of the *rel*_*Mtb*_ gene with the immature dendritic cell-targeting gene encoding chemokine MIP-3α/CCL20. To augment mucosal immune responses, intranasal delivery was also evaluated. We found that the intramuscular *MIP-3α*/*rel*_*Mtb*_ (fusion) vaccine potentiates isoniazid activity more than a similar DNA vaccine expressing *rel*_*Mtb*_ alone in a chronic TB mouse model (absolute reduction of *Mtb* burden: 0.63 log_10_ colony-forming units, P=0.0001), inducing pronounced *Mtb*-protective immune signatures. The intranasal fusion vaccine, an approach combining *rel*_*Mtb*_ fusion to MIP-3α and intranasal delivery, demonstrated the greatest therapeutic effect compared to each approach alone, as evidenced by robust Th1 and Th17 responses systemically and locally and the greatest mycobactericidal activity when combined with isoniazid (absolute reduction of *Mtb* burden: 1.13 log_10_, P<0.0001, when compared to the intramuscular vaccine targeting *rel*_*Mtb*_ alone). This DNA vaccination strategy may be a promising adjunctive approach combined with standard therapy to shorten curative TB treatment, and also serve as proof-of-concept for treating other chronic infections.

## Introduction

Tuberculosis (TB) is a major cause of morbidity, and the second leading infectious killer after COVID-19 worldwide (1). The current six-month regimen, consisting of isoniazid (INH), rifampin, pyrazinamide and ethambutol, has high efficacy against drug-sensitive TB, but its length and complexity contributes to treatment interruptions that jeopardize cure and promote drug resistance (2, 3). Although novel, treatment-shortening antibiotic regimens have shown promising results in international clinical trials (4, 5), the resources required for direct observation of daily treatment and the associated costs may still pose barriers to their implementation in TB-endemic countries. Recent work has focused on adjunctive, host-directed strategies to simplify and shorten the course of TB therapy (6).

The need for prolonged TB treatment is believed to reflect the unique ability of a subpopulation of *Mycobacterium tuberculosis* (*Mtb*) bacilli within the infected host to remain in a nonreplicating, persistent state (7) characterized by tolerance to first-line anti-TB drugs, like INH, which more effectively targets actively dividing bacilli (8-11). One of the key bacterial pathways implicated in antibiotic tolerance is the stringent response, which is regulated by the (p)ppGpp synthase/hydrolase, Rel_Mtb_ (12, 13). Rel_Mtb_ deficiency results in defective *Mtb* survival under nutrient starvation (14), in mouse lungs (15) and mouse hypoxic granulomas (16), reduced virulence in guinea pigs (17) and C3HeB/FeJ mice (18), and increased susceptibility of *Mtb* to INH in mouse lungs (18), rendering Rel_*Mtb*_ an attractive target for novel anitubercular therapies, including for drug-resistant TB (13).

We previously showed that intramuscular (IM) delivery of a DNA vaccine expressing rel_*Mtb*_ enhanced the mycobactericidal activity of INH in a murine TB model (3, 19). In independent studies, our group has shown the enhanced efficacy of vaccines when the antigen of interest was fused to the gene encoding the chemokine Macrophage Inflammatory Protein-3 alpha/C-C Motif Chemokine Ligand 20 (MIP-3α/CCL20) (20-22). This chemokine targets the antigen of interest to immature dendritic cells (DCs) and, compared to vaccines without the MIP-3α component, has shown enhanced T-cell responses in both melanoma and malaria model systems (20-22). Since T-cell immunity is required to control *Mtb* infection (6), we hypothesized that fusion of *rel*_*Mtb*_ to the chemokine gene MIP-3α (*MIP-3α/rel*_*Mtb*_ or “fusion vaccine”) would enhance the immunogenicity of the *rel*_*Mtb*_ vaccine and further potentiate the mycobactericidal activity of INH *in vivo*.

Protection against pulmonary TB is associated with the ability of anti-*Mtb* T-cells to exit the pulmonary vasculature and enter into the lung parenchyma and airways (23). Intranasal (IN) vaccination has been shown to promote recruitment of antigen-experienced T cells to these restricted lung compartments in contrast to parenteral immunization (23). Thus, we also hypothesized that IN administration of the vaccine expressing *rel*_*Mtb*_ alone or the fusion vaccine would further augment T-cell responses within the lung, the primary site of *Mtb* infection.

Here, we present our bacteriological and immunological findings on IM vs. IN immunization with a DNA vaccine expressing *rel*_*Mtb*_ alone or the fusion construct in a murine model of chronic TB. Our results indicate that *rel*_*Mtb*_ fusion to *MIP-3α* and IN vaccination (“optimized vaccination strategy”) yielded the highest additive therapeutic effect compared to each single approach alone.

## Results

### MIP-3α fusion and IN delivery of the *rel*_*Mtb*_ vaccine individually increase the mycobactericidal activity of INH in a murine model of chronic TB

Four weeks after *Mtb* aerosol infection, C57BL/6 mice were treated daily with human-equivalent doses of oral INH for 10 weeks (Figure 1B). The *rel*_*Mtb*_ or the fusion vaccine [Figure 1A, detailed sequences are available in the Supplementary Appendix] was administered via the IM [Standard Dose (SD) or High Dose (HD)] or IN (HD) routes weekly for 3 weeks. The original SD *rel*_*Mtb*_ vaccine given IM, which previously demonstrated therapeutic adjunctive activity together with INH (3, 19), served as our baseline comparator in this study (“comparator” vaccine). For IN delivery, only the HD vaccine was tested to compensate for the anticipated reduced plasmid uptake without electroporation, which cannot be used with this vaccination route. One negative control group received no treatment, while another group received INH only. All vaccinated groups received INH in addition to the tested vaccines. Since DNA vaccination alone did not exhibit significant mycobactericidal activity in prior work (19), this group was not included in the present study.

**Figure 1.**
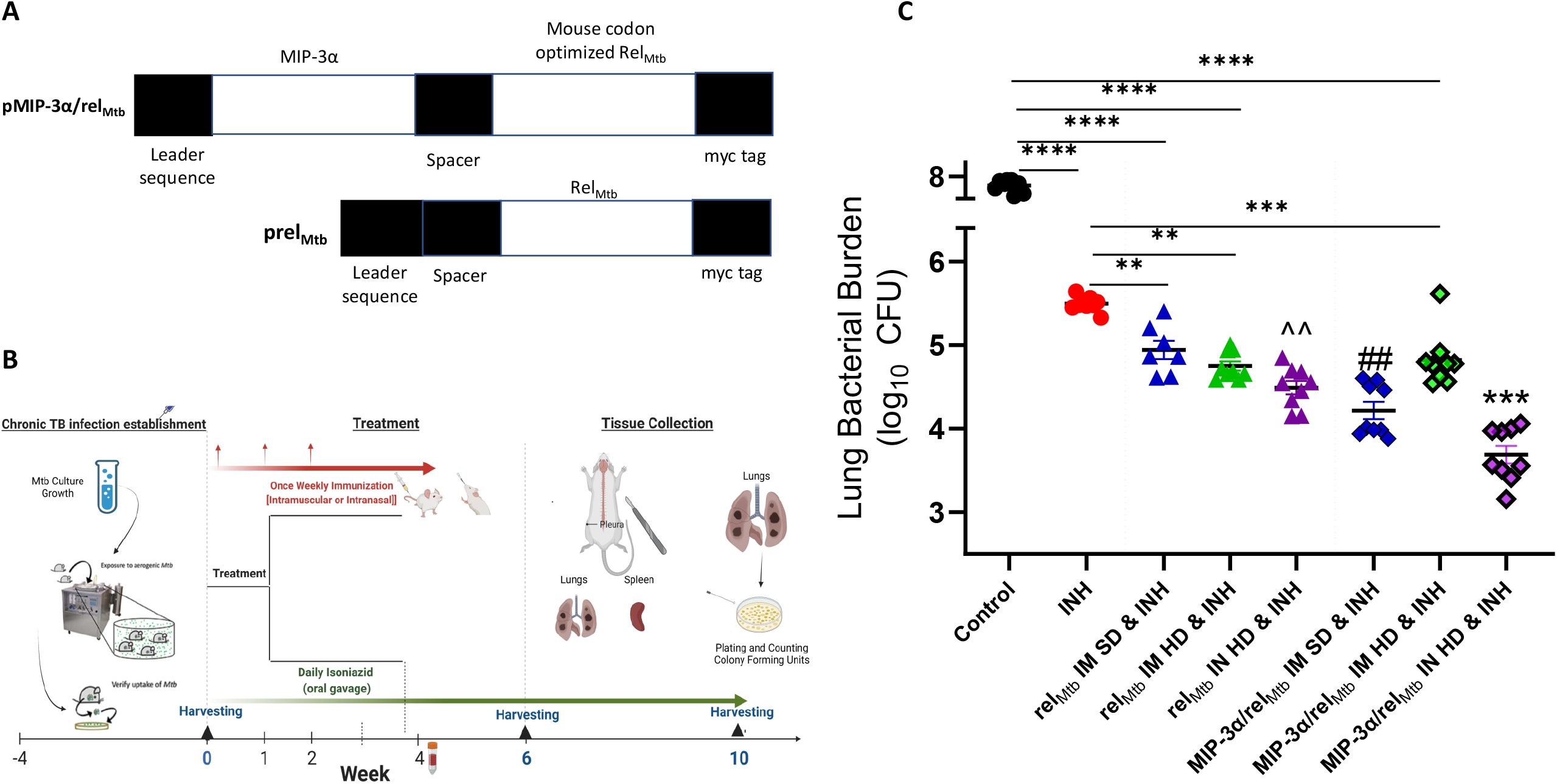
MIP-3α fusion and IN delivery of vaccine expressing relMtb increase the mycobactericidal activity of INH in a murine model of chronic TB. (A) Diagrammatic representation of the MIP-3α/relMtb and relMtb DNA constructs used for immunization. (B) Timeline of the Mtb challenge study, (C) Scatterplot of lung mycobacterial burden at 10 weeks after the primary vaccination per vaccination group: IN delivery of a DNA vaccine expressing *rel*_*Mtb*_ or IM delivery of a DNA vaccine expressing *MIP-3α/rel*_*Mtb*_ enhances the mycobactericidal activity of INH *in vivo* compared to IM delivery of *rel*_*Mtb*_ vaccine. The greatest therapeutic effect is demonstrated after IN *MIP-3*^*α*^*/rel*_*Mtb*_ vaccination, which is more efficacious compared to any other group. *Mtb*: *Mycobacterium tuberculosis*, IM: Intramuscular, IN: Intranasal, SD: Standard Dose, HD: High Dose, CFU: colony-forming units, INH: Isoniazid. All statistically significant P values are available in Supplementary Table 1. *** = significant difference from all the experimental groups (at least P< 0.001). ## = significant difference from all the experimental groups except *rel*_*Mtb*_ IN (at least P< 0.01). ^^ = significant difference from all except *MIP-3α/rel*_*Mtb*_ IM (at least P< 0.01).

At 10 weeks after primary vaccination by the IM route, greater potentiation of mycobactericidal activity of INH was observed in groups receiving the SD fusion vaccine compared to mice receiving the comparator vaccine [absolute reduction of mycobacterial burden: 0.63 log_10_ colony-forming units (P=0.0001)]. Also, IN vaccination with the *rel*_*Mtb*_ vaccine significantly enhanced the mycobactericidal activity of INH compared to IM vaccination with the comparator vaccine [absolute reduction of mycobacterial burden: 0.52 log_10_ colony-forming units (P=0.0052), Figure 1C]. Interestingly, IM vaccination with the HD fusion or *rel*_*Mtb*_ vaccines did not significantly increase the mycobactericidal activity of INH relative to the comparator vaccine, indicating that the improved therapeutic efficacy observed with the IN vaccination approach is not simply dose-related. (Figure 1C).

IN vaccination with the fusion vaccine (hereafter referred to as “optimized vaccination strategy”) showed the greatest additive therapeutic effect in combination with INH compared to any other experimental group; there was an absolute reduction in lung bacillary load of 1.13 log_10_ relative to the comparator vaccine, of 0.5 log_10_ relative to the IM-delivered SD fusion vaccine (P=0.0058; Figure 1C), and of 0.61 log_10_ relative to the IN-delivered *rel*_*Mtb*_ vaccine (P<0.0001; Figure 1C). At 10 weeks post-primary vaccination, the optimized vaccination strategy resulted in the greatest reduction in normalized mean lung weight, which serves as a proxy for totatl lung inflammation, relative to the INH only group (relative reduction in normalized lung weight by 42.4%; P<0.0002) and untreated control group (relative reduction in normalized lung weight by 66.3%; P<0.0001) (Supplementary Figure 1A). All the individual comparisons between the different experimental groups are available in Supplementary Table 1. Mean lung mycobacterial burdens at implantation (−4 weeks), initiation of treatment (0 weeks), and at 6 weeks and 10 weeks after the initiation of treatment for each experimental group are shown in Supplementary Figure 1B. Gross pathology photographs of representative lungs per experimental group are available in Supplementary Figure 1C.

### IM vaccination with the fusion vaccine elicits a robust systemic Th1 response

For simplicity, we have included only the SD IM vaccine groups and not the HD IM groups, since both IM groups yielded similar microbiological outcomes. Relative to the comparator vaccine, the IM fusion vaccine elicited substantially higher numbers of Rel_Mtb_-specific, IFN-γ-producing CD4+ and CD8+ T lymphocytes in the spleens (P<0.0001 and P<0.0001; Figure 2A and B), but not in the lungs of infected mice (Figure 2C and 2D). Relative to the comparator vaccine, the IM-delivered fusion vaccine was also associated with significantly increased numbers of Rel_Mtb_-specific, TNF-α-producing CD4+ T cells (P=0.0076, Figure 2E) but not CD8+ T cells (Figure 2F) in the spleens of *Mtb-*infected mice. Significant production of Rel_Mtb_-specific, IL-2-producing CD4+ and CD8+ T cells in the spleens of infected mice was also noted after IM vaccination with the fusion vaccine relative to the comparator vaccine (P=0.005 and P<0.0001, respectively, Figures 2G and 2H). IM vaccination with the fusion vaccine or the comparator vaccine elicited similar numbers of Rel_Mtb_-specific, IL-17α-producing CD4+ T cells in infected murine lungs or spleens (Figure 2I and 2J). All the individual statistical comparisons of IM vaccination with the fusion vaccine relative to all the other experimental groups are available in Supplementary Table 2.

**Figure 2.**
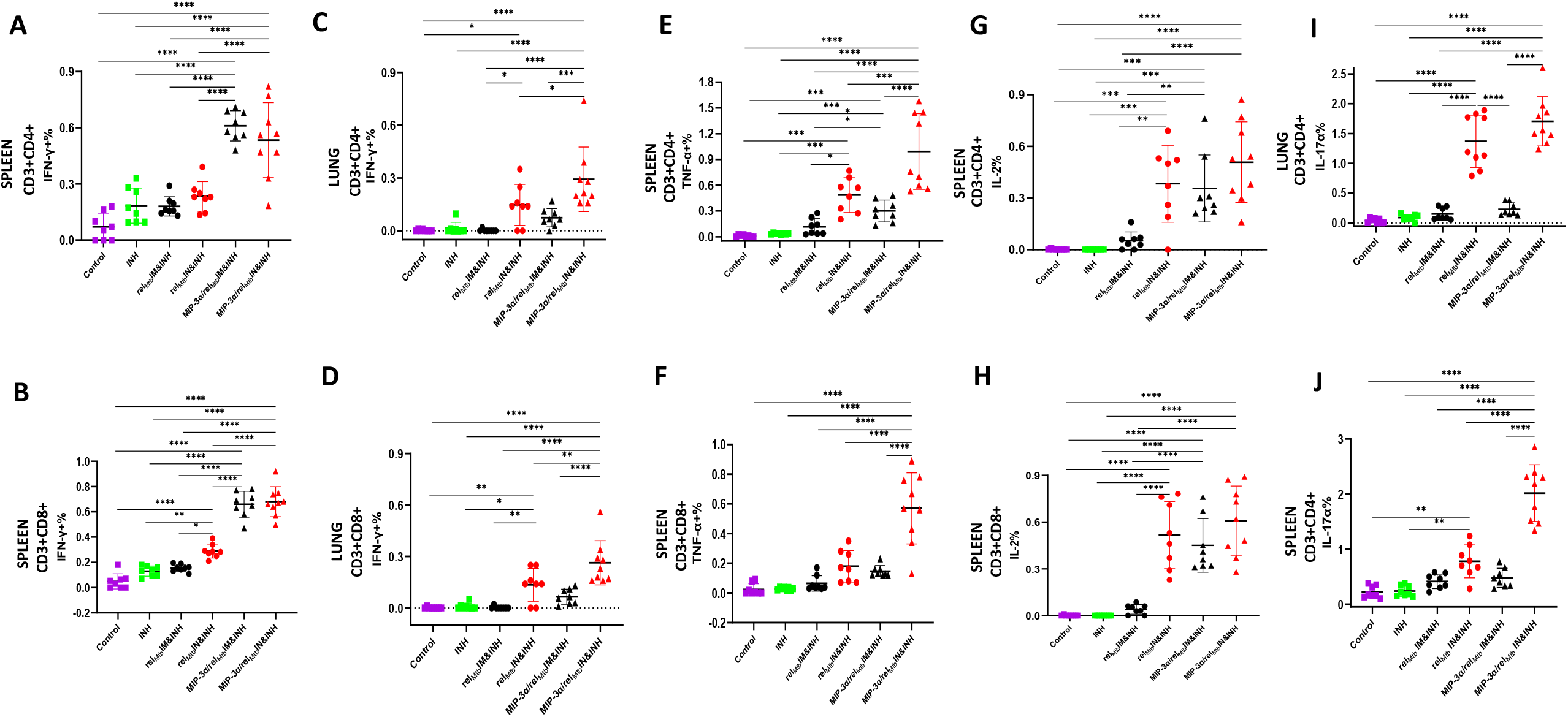
T-cell responses in murine tissues 6 weeks after *Mtb* challenge: IN vaccination with *rel*_*Mtb*_ or IM vaccination with *MIP-3α*/*rel*_*Mtb*_ elicits higher Th17 or Th1 response compared to IM vaccination with rel_*Mtb*_,while IN vaccination with *MIP-3α/rel*_*Mtb*_ offers the most robust systemic and local Th1 and Th17 responses of all experimental groups. Rel_Mtb_-specific IFN-γ^-^producing CD4+ T cells (A) and CD8+ T cells (B) in spleens; Rel_Mtb_-specific IFN-γ^-^producing CD4+ T cells (C) and CD8+ T cells (D) in lungs; Rel_Mtb_-specific, TNF-α^-^producing CD4+ T cells (E) and CD8+ T cells (F) in spleens; Rel_Mtb_-specific TNF-α^-^producing CD4+ T cells (G) and CD8+ T cells (H) in lungs; Rel_Mtb_-specific, IL-17α^-^producing CD4+ T cells in lungs (I) and spleens (J) (flow cytometry-intracellular staining).. IM: Intramuscular (standard dose is shown), IN: Intranasal (high dose is shown). All statistically significant P values are available in Supplementary Table 2. Y-axis scales are different among cytokines and between tissues in order to better demonstrate differences between groups where cytokine expression levels were lower.

In an independent immunogenicity study using uninfected animals (Figure 3A), we also tested the T-cell responses in additional murine tissues, including draining lymph nodes (LNs) (Figure 3B) and peripheral blood mononuclear cells (PBMCs) (Figure 3C and 3D), in addition to spleens and lungs (Supplementary Figure 2). Relative to the comparator, IM vaccination with the fusion vaccine elicited more Rel_*Mtb*_-specific, IL-17α-producing CD4+ T cells in the draining LNs and PBMCs (P=0.036 and P=0.036 respectively, Figure 3B and 3C). Similarly, IM delivery of the fusion vaccine elicited more Rel_*Mtb*_-specific, TNF-α-producing CD4+ T cells in the PBMC population starting at 28 days (P=0.036) and peaking at 42 days (P=0.036) after primary vaccination (Figures 3D). All the individual statistical comparisons of IM-delivered fusion vaccine with all the other experimental groups in uninfected animals are available in Supplementary Table 3.

**Figure 3.**
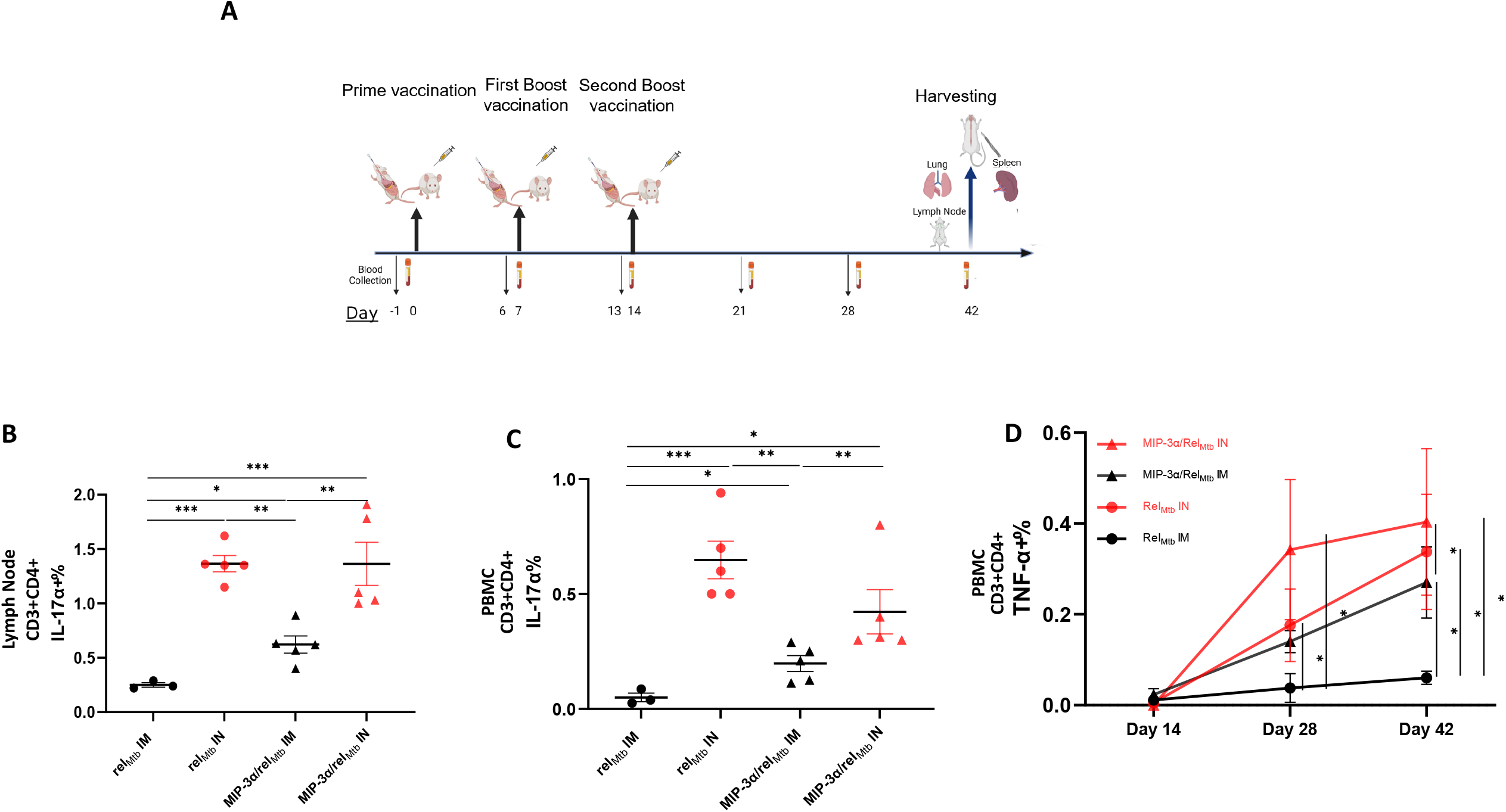
T-cell responses in non-infected murine tissues 6 weeks after prime vaccination: IN vaccination with rel_*Mtb*_ *or MIP-3α/rel*_*Mtb*_ and IM vaccination with *MIP-3α/rel*_*Mtb*_ elicited stronger IL-17α CD4+ T-cell responses in cells extracted from draining LNs, PBMCs and higher TNF-α CD4+ T cells in PBMCs compared to IM vaccination with *rel*_*Mtb*_. (A) Timeline of the immunogenicity study. Rel_Mtb_-specific, IL-17α^-^ producing CD4+ T cells in draining LNs (B) and PBMCs (C) and Rel_Mtb_-specific, TNF-α-producing CD4+ T cells (D) in PBMCs. (flow cytometry-intracellular staining). All statistically significant P values are available in Supplementary Table 3. IM: Intramuscular, IN: Intranasal, LNs: Lymph nodes, PBMCs: Peripheral Blood Mononuclear Cells. Y-axis scales are different among cytokines and between tissues in order to better demonstrate differences between groups where cytokine expression levels were lower.

### IN vaccination with *MIP-3α/rel*_*Mtb*_, the optimized vaccination strategy, elicits the most robust systemic and local Th1/Th17 responses relative to any other vaccination strategy

Having established that IM vaccination with the fusion vaccine induces enhanced systemic Th1 responses relative to the comparator, we next assessed the effect of the IN vaccination route on immune responses elicited by the fusion vaccine. The optimized vaccination strategy group had greater numbers of Rel_*Mtb*_-specific, IFN-γ-producing CD4+ and CD8+ T cells in the lungs of *Mtb-*infected mice (P=0.003 and P<0.0001, Figure 2C and 2D, respectively) compared to the IM-delivered fusion vaccine. The optimized vaccination strategy also induced higher numbers of Rel_*Mtb*_-specific, TNF-α-producing CD4+ and CD8+ Tcells in the spleens compared to the IM-delivered fusion vaccine (P<0.0001 and P<0.0001, Figure 2E and 2F, respectively). Rel_*Mtb*_-specific, IL-2-producing CD4+ and CD8+ T cells in the spleens of infected mice were similarly high irrespective of the route of delivery of the fusion vaccine (Figure 2G and 2H). Infected mice receiving the optimized vaccination strategy showed a significantly higher percentage of Rel_*Mtb*_-specific, IL-17α-producing CD4+ T cells in the lungs and spleens compared to those receiving the fusion vaccine by the IM route (P<0.0001 and P<0.0001, Figure 2I and 2J).

Relative to IN vaccination with the *rel*_*Mtb*_ vaccine, the optimized vaccination strategy elicited greater numbers of Rel_*Mtb*_-specific, IFN-γ-producing CD4+ and CD8+ T cells in the spleens (P<0.0001 and P<0.0001, Figure 2A and 2B, respectively) and in the lungs (P=0.034 and P=0.0064, Figure 2C and 2D, respectively) of infected mice. The optimized vaccination strategy also resulted in higher numbers of Rel_*Mtb*_-specific, TNF-α-producing CD4+ and CD8+ T cels in the spleens compared to mice receiving the *rel*_*Mtb*_ vaccine (P=0.0003 and P<0.0001, Figure 2E and 2F, respectively). Rel_*Mtb*_-specific IL-2-producing CD4+ and CD8+ T lymphocytes in the spleens were similarly high in the optimized vaccination group and the IN *rel*_*Mtb*_ group (Figure 2G and 2H). Mice receiving the optimized vaccination strategy showed a significantly higher percentage of Rel_*Mtb*_-specific, IL-17α-producing CD4+ T cells in the spleens compared to mice receiving the *rel*_*Mtb*_ vaccine by the IN route (P<0.0001, Figure 2J), but no difference between the two groups was noted in the lungs (Figure 2I). Of note, no IL-17α response was observed at these sites following IM immunization.

Focusing on the differences in the immune responses in the IN *rel*_*Mtb*_ and the comparator groups, the former approach induced a significantly increased proportion of Rel_*Mtb*_-specific, IFN-γ-producing CD4+ and CD8+ T cells in the lungs (P=0.044 and P=0.006; Figure 2C and D), and of IFN-γ-producing CD8+ T cells in the spleens of infected mice (P=0.012, Figure 2B). IN vaccination with the *rel*_*Mtb*_ vaccine also induced significantly higher numbers of Rel_Mtb_-specific, TNF-α-producing CD4+ T cells in the spleens relative to the comparator group (P=0.017, Figure 2C). Moreover, relative to the comparator, IN vaccination with the *rel*_*Mtb*_ vaccine significantly increased Rel_Mtb_-specific, IL-2-producing CD4+ and CD8+ T cells in the spleens [P=0.0018 and P<0.001,Figures 2G and H), and a higher percentage of Rel_Mtb_-specific, IL-17α-producing CD4+ T cells in the lungs (P<0.0001; Figure 2I), but not in the spleens (Figure 2J). All the individual statistical comparisons between experimental groups are available in Supplementary Table 2.

In an independent immunogenicity study using uninfected animals (Figure 3A), we found that, compared to fusion vaccine given IM, the optimized vaccination strategy elicited more robust Rel_*Mtb*_-specific, IL-17α CD4+ T-cell responses in the draining LNs (P=0.0033, Figures 3B) and higher Rel_*Mtb*_-specific, IL-17α CD4+ T-cell responses in PBMCs (P=0.0079, Figure 3C). The optimized vaccination strategy also induced a higher percentage of Rel_*Mtb*_-specific, TNF-α-producing CD4+ T cells at 42 days compared to the fusion vaccine given IM (P=0.03, Figure 3D). IN vaccination with the *rel*_*Mtb*_ vaccine elicited a higher percentage of Rel_Mtb_-specific, IL-17α-producing CD4+ T cells relative to the comparator in the draining LNs (P=00003, Figure 3B) and PBMCs (P=0.0007, Figure 3C). Finally, by day 42 after primary vaccination, IN delivery of the *rel*_*Mtb*_ vaccine induced higher proportions of Rel_*Mtb*_-specific, TNF-α-producing CD4+ T cells in PBMCs relative to the comparator (P=0.036, Figure 3D). The individual statistical comparisons of immunological outcomes in uninfected mice resulting from different vaccination protocols are available in Supplementary Table 3.

Collectively, the optimized vaccination strategy, which was shown to have the most favorable microbiological outcomes over any other tested vaccination approache (Figure 1C), was also found, in parallel, to significantly increase the production of multiple cytokines associated with *Mtb* control, both systemically and at the site of infection. More specifically, compared to any other approach, the optimized vaccination strategy induced the highest numbers of Rel_*Mtb*_-specific CD4+ and CD8+ T cells producing IL17-α,TNF-α, IFN-γ, and IL-in the spleens and lungs of *Mtb*-infected animals (Figure 4A, 3B, 3C, Supplementary Table 2), as well as the highest normalized production of each individual cytokine (Figures 4D, 4E, 4F, 4G, 4H, 4I).

**Figure 4.**
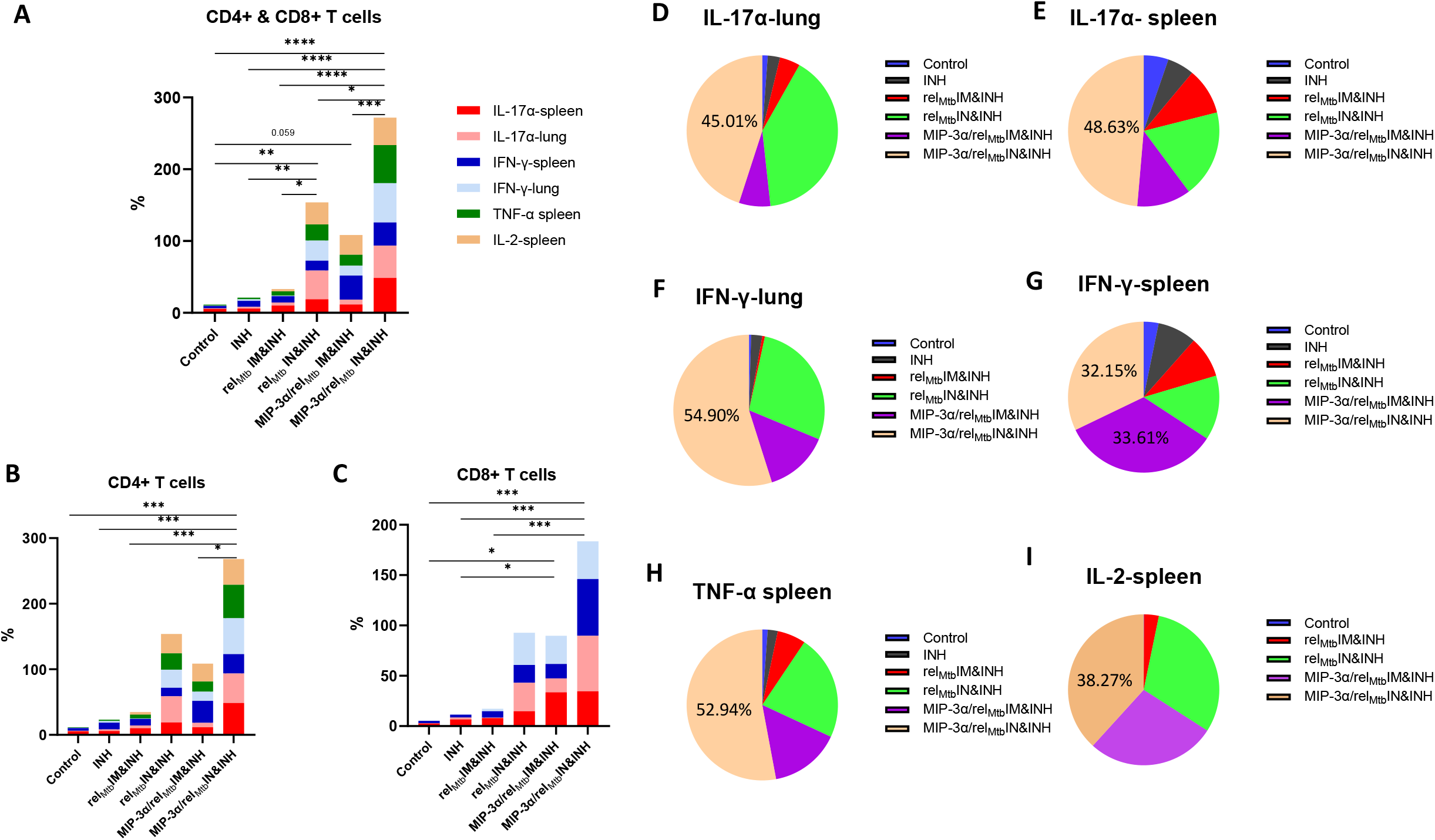
IN vaccination with *MIP-3α/rel*_*Mtb*_ increases the simultaneous production of multiple cytokines associated with *Mtb* control, systemically and at the site of infection. IN *MIP-3α/rel*_*Mtb*_ vaccination group was found to have the highest aggregate production of the IL17-α,TNF-α, IFN-γ, and IL-2-producing CD4+ and CD8+ T cells in the spleens and lungs of *Mtb*-infected animals compared to any other group (A, B, C), but also had the highest normalized production of each cytokine individually across experimental groups (D, E, F, G, H, I).

## Discussion

The development of novel immunotherapeutic regimens that synergize with antibiotics to accelerate curative TB treatment is an attractive strategy for improving medical adherence and treatment completion rates, and for reducing costs (24). In the current study, we show that *MIP-3α* fusion and the IN route of delivery individually enhance the therapeutic adjunctive activity of a DNA vaccine targeting an *Mtb* persistence antigen in a murine model of chronic TB. Importantly, the combined approach, i.e., IN immunization with a DNA fusion vaccine expressing *MIP-3α/rel*_*Mtb*_, was accompanied by additive Th1/Th17 responses, both systemically and at the site of infection. This novel optimized vaccination strategy may be a promising adjunctive therapeutic approach in combination with standard anti-TB therapy.

A true functional immunological signature to predict adequate TB control is still lacking, but it is clear that CD4+ and CD8+ T cells are critical in developing immunity against *Mtb* (25-29). T-cell immunity to TB is likely mediated by a variety of T cells, especially those mediating Th1 and Th1/Th17-like responses (30). Chronic antigenic stimulation drives antigen-specific CD4+ T-cell functional exhaustion during murine *Mtb* infection (31), with important implications for TB vaccine design. Thus, subdominant *Mtb* antigens during chronic *Mtb* infection, including Rel_Mtb,_ which is induced during antitubercular treatment (3), may represent promising targets for therapeutic vaccines in an effort to “re-educate” the immune system to tailor host anti-TB responses. In the current study, we focused on improving Rel_Mtb-_specific T-cell responses by enhancing the engagement of immature DCs.

Immature DCs are critical for the activation of adaptive immunity, and, eventually, mature DCs trigger antigen-specific naïve T cells (32). Of note, only a small minority of DCs are attracted to sites of immunization (32), and, in the case of HIV and TB infections, a proportion of the attracted DCs may be dysfunctional (33). Fusion of the antigen of interest to the chemokine MIP-3α (or CCL20) targets the antigen to immature DCs (24). It has been shown that following naked DNA vaccination, epidermal cells secrete the antigen of interest-chemokine MIP-3α fusion construct (20, 34). The secreted fusion construct is taken up and internalized by skin Langerhans cells via the receptor for this chemokine, which is called CCR6 (34). The complex is then processed and presented in draining LNs to elicit efficient cellular and humoral responses (34). Enhanced efficacy has been shown compared to antigen-only vaccines in various systems (20-22). In a mouse melanoma model, our group has demonstrated that IM immunization with a DNA vaccine containing a fusion of *MIP-3α* with the tumor antigen gene *gp100/Trp2* elicited greater numbers of tumor antigen-specific T cells and offered greater therapeutic benefit compared to the cognate vaccine lacking the *MIP-3α* fusion (20, 21). Importantly, MIP-3α has also been shown to play a key role in driving DC recruitment to the nasal mucosa (35). Indeed, we found that IM vaccination with the *MIP-3α* fusion construct conferred increased antigen-specific systemic Th1 responses (IFN-γ, TNF-α, Il-2 in the spleens and TNF-α in PBMCs), but also Th17 responses in the draining LNs and PBMCs, relative to the *rel*_*Mtb*_ construct alone. Interestingly, no Th1 or Th17 responses were observed in the lungs, the primary site of the infection. This fusion vaccination strategy, i.e., IM vaccination with *MIP-3α/rel*_*Mtb*_, yielded improved microbiological outcomes when combined with INH compared to the non-fused *rel*_*Mtb*_ vaccine.

Compelling evidence suggests that protection against respiratory pathogens, such as *Mtb*, is dependent on the presence of pathogen-specific immune cells at the primary site of infection (24, 36, 37). Pre-clinical studies have shown that parenteral immunization with TB vaccines can drive robust antigen-specific T-cell responses in the periphery, but these cells are unable to rapidly enter the restricted lung mucosal compartments and largely fail to restrict *Mtb* replication (38). In contrast, respiratory mucosal immunization generates a long-lasting population of tissue-resident T cells expressing homing molecules to allow preferential migration and residence in the airway lumen and lung parenchyma (37-39). These immune cells generate a ‘first line of defense’ by establishing pathogen-specific immunity at the site of entry, providing markedly enhanced control against pulmonary *Mtb* infection (37-39). Importantly, after IN vaccination, these antigen-experienced, lung-resident, T-cells have been shown to produce IL-17α in addition to IFN-γ, expanding the known signature panel that may confer enhanced TB immunity (40-43). In the current study, we have shown for the first time that IN delivery of a vaccine (*rel*_*Mtb*_) enhances the bactericidal activity of an antitubercular drug (INH) relative to IM delivery of the same vaccine. IN immunization resulted in more robust Th1 (IFN-γ, TNF-α, IL-2) and Th17 responses (IL-17α) systemically, but also in the lungs (IFN-γ and IL-17), the primary site of infection. Importantly, the combined approach of *MIP-3α* fusion and the IN route of immunization yielded the greatest additive adjunctive mycobactericidal activity with INH in murine lungs, resulting in an approximate 100-fold reduction in lung bacterial burden compared to INH alone. This immunization protocol was accompanied by the most robust, additive Th1 and Th17 responses, both systemically and in the lungs of *Mtb*-infected mice.

In conclusion, we have shown that IN immunization with a DNA vaccine expressing *MIP-3α/rel*_*Mtb*_ generates strong, additive Th1 and Th17 responses and significantly potentiates the mycobactericidal activity of the first-line drug, INH. Further studies are required to elucidate the relative importance of the different effector mechanisms elicited by this immunization strategy and to refine our understanding of the host-pathogen interactions that result in the improved therapeutic effects. Ultimately, the potential utility of this vaccination combination strategy must be evaluated as an adjunctive therapeutic intervention in shortening the duration of curative treatment for active TB.

## Methods

### Bacteria and growth conditions

Wild-type *Mtb* H37Rv was grown in Middlebrook 7H9 broth (Difco, Sparks, MD) supplemented with 10% oleic acid-albumindextrose-catalase (OADC, Difco), 0.2% glycerol, and 0.05% Tween-80 at 37°C in a roller bottle (3).

### Antigen preparation

The previously generated *rel*_*Mtb*_ expression plasmid, pET15b[rel_Mtb_] (3, 19), was used for expression and purification of recombinant Rel_Mtb_ protein. *Escherichia coli* BL21 (DE3) RP competent cells (Stratagene) were transformed with pET15b[rel_Mtb_]. Transformed bacteria were selected with ampicillin (100 mg/ml), and cloning was confirmed by DNA sequencing. Protein expression was performed using standard protocols and purification was performed using Ni-NTA Agarose (Qiagen). Recombinant Rel_*Mtb*_ protein (87 kDa) was purified from the cell lysate using a Ni-NTA resin column. The purity was confirmed by SDS-PAGE gel and immunoblot analyses. The protein concentration was determined using a BCA protein assay with BSA as the standard (Thermo Fisher). Recombinant Rel_Mtb_ has been shown previously to retain (p)ppGpp synthesis and hydrolysis activities and can serve as an antigen to measure Rel_Mtb_-specific T-cell responses *ex vivo* (3, 19).

### DNA vaccines

The plasmid pSectag2B encoding the full-length *rel*_*Mtb*_ gene was used as the *rel*_*Mtb*_ DNA vaccine (19). The *rel*_*Mtb*_ gene was codon-optimized (Genscript) and fused to the mouse *MIP-3α* gene. The fusion product was cloned into pSectag2B, serving as the *MIP-3α/rel*_*Mtb*_ DNA, fusion, vaccine or (Figure 1A, detailed sequence in Supplementary Appendix). Proper insertion was confirmed by sequencing and the expression of target genes was confirmed by transfection of 293T cells in lysates and supernatants. Vaccination plasmids were selected by ampicillin (100 μg/ml) and extracted from *E. coli* DH5-α (Invitrogen™ ThermoFisher Scientific, Waltham, MA) using Qiagen® (Germantown, MD) EndoFree® Plasmid Kits and were diluted with endotoxin-free 1xPBS.

### *Mtb* challenge study in mice

Male and female C57BL/6 mice (8-10-week-old, The Jackson Laboratory) were aerosol-infected with ∼100 bacilli of wild-type *Mtb* H37Rv using a Glas-Col Inhalation Exposure System (Terre Haute, IN). After 28 days of infection, the mice received INH 10 mg/kg dissolved in 100 ml of distilled water by esophageal gavage once daily (5 days/week) and were randomized to receive *rel*_*Mtb*_ or the fusion vaccine by the intramuscular (standard dose: 20 μg or high dose: 200 μg) or intranasal (high dose: 200 μg) route. The mice were vaccinated three times at one-week intervals. Each plasmid was delivered IM or IN after mice were adequately anesthetized by vaporized isoflurane. For IM immunizations, each plasmid was injected bilaterally into the quadriceps femoris muscle of the mice (50 μL in each quadriceps), followed by local electroporation using an ECM830 square wave electroporation system (BTX Harvard Apparatus Company, Holliston, MA, USA). Each of the two-needle array electrodes delivered 15 pulses of 72 V (a 20-ms pulse duration at 200-ms intervals) (19). For IN immunizations, each plasmid was administered into both nostrils (50 μL in each nostril) and mice were monitored in the upright position until complete recovery and vaccine absorption were assured. The mice were sacrificed 6 weeks and 10 weeks after treatment initiation. The spleens and left lungs were harvested and processed into single-cell suspensions. The right lungs were homogenized using glass homogenizers. Serial tenfold dilutions of lung homogenates in PBS were plated on 7H11 selective agar (BD) at the indicated time points. Plates were incubated at 37° C and colony-forming units (CFU) were counted 4 weeks later (3, 19).

### Immunogenicity studies in mice

Male and female C57BL/6 mice (8-10-week-old, Charles River Laboratory) were randomized to receive the *rel*_*Mtb*_ or the fusion DNA vaccine by the IM (20 or 200 μg) or IN (200 μg) route. The mice were sacrificed 6 weeks after the primary vaccination. Spleens, draining LNs, lungs and PBMCs were collected and processed into single-cell suspensions.

### Intracellular Cytokine Staining, Flow Cytometry Analysis and FluoroSpot

All the single-cell suspensions from spleens, draining LNs, lungs and PBMCs were stimulated individually with purified recombinant Rel_Mtb_ protein at 37°C (3, 19) for various time intervals, from 12 hrs (IFN-γ, IL-17α, IL-2) to 24 hrs (TNF-α), depending on the cytokine of interest. For Intracellular Cytokine Staining (ICS), GolgiPlug cocktail (BD Pharmingen, San Diego, CA) was added for an additional 4 hours after stimulation (total, 16 and 28 hrs, respectively) and cells were collected using FACS buffer (PBS + 0.5% Bovine serum albumin (Sigma-Aldrich, St. Louis, MO), stained with Zombie NIR™ Fixable Viability Kit (Biolegend Cat. No.: 423105) for 30 min, washed with PBS buffer, surface proteins were stained for 20 min, cells were fixed and permeabilized with buffers from Biolegend intracellular fixation/ permeabilization set following manufacturer protocols (Cat. No. 421002), intracellular proteins were stained for 20 min, and samples were washed and resuspended with FACS buffer. The following anti-mouse mAbs were used for ICS: PercPCy5.5 conjugated anti-CD3 (Biolegend Cat. No 100217), FITC conjugated anti-CD4 (Biolegend Cat. No 100405), Alexa700 conjugated anti-CD8 (Biolegend Cat. No. 155022), PECy7 conjugated anti-TNF-α, (Biolegend Cat. No. 506323), APC conjugated anti-IFN-γ, (Biolegend Cat. No. 505809), BV421 conjugated anti-IL-2, (Biolegend Cat. No 503825), PE conjugated anti-IL-17α (Biolegend Cat. No 506903). The Attune™ NxT (Thermo Fisher Scientific, Waltham, MA), and a BD™ LSRII flow cytometer was used. Flow data were analyzed by FlowJo Software (FlowJo 10.8.1, LLC Ashland, OR). Flow analysis included alive, gated, total T lymphocytes, including CD4+ T- and CD8+ T-cell subpopulations. For simplicity, the CD8+ subpopulation analysis is not reported if no substantial population to allow comparisons was detected (e.g., IL-17α). For FluoroSpot assays (Supplementary data), kits with pre-coated plates for enumeration of cells secreting IFN-γ and IL-17A were purchased from Mabtech (Cat. No. FSP-414443-2). Spots were enumerated on an AID iSpot EliSpot/ FluoroSpot Reader.

### Statistics

Pairwise comparisons of group mean values for log_10_ CFU (microbiology data) and flow cytometry data were made by using one-way analysis of variance followed by Tukey’s multiple comparisons test. Prism 9.3 (GraphPad Software, Inc. San Diego, CA) was utilized for statistical analyses and figure creation. For correlation of two nominal predictor variables to a continuous outcome variable, two-way analysis of variance followed by Tukey’s multiple comparisons test was utilized. To illustrate the aggragate cytokine data per group (Figure 4), Fraction of total analysis was used and is displayed in stacked bars and pie charts. In Figure 4, cytokine data were normalized (the total sum of all the experimental groups= 100%, total absolute number of alive cells=30,000). All error bars represent the estimation of the standard error of the mean, and all midlines represent the group mean. A significance level of α ≤ 0.05 was set for all experiments.

### Study Approval

All procedures were performed according to protocols approved by the Johns Hopkins University Institutional Animal Care and Use Committee.

## Acknowledgements

This work was supported by NIH grants: R21AI140860 and R01 AI148710 to RBM and PCK and T32 AI007291 to SK. The content is solely the responsibility of the authors and does not necessarily represent the official views of the National Institutes of Health. Illustrations Figure 1B and, 3A were created with BioRender.com.

## Author Contributions

SK, RBM and PCK conceived and designed the study and wrote the manuscript. JTG and SK performed the fusion vaccine construction. SK performed transfection of 293T cells and ran western blots in lysates and supernatants. SK and KC performed Rel_Mtb_ protein expression, purification and verification. SK performed growth, selection and extraction of vaccination plasmids. SK and TK performed the *in vivo* vaccinations (immunogenicity and challenge study). SK, PN and TK administered daily INH treatments to mice (challenge study). SK performed survival mouse tail bleeds weekly and isolated PBMCs for further analysis (immunogenity study). SK, PN, TK, JTG, JRC, DQ, KC, AS, YH, SKA, and CD performed animal harvesting and subsequent animal tissue processing (immunogenicity and challenge study). PN, RT and SC contributed to the *in vivo* challenge study design. SK, TK and PN plated the homogenates of the infected lungs and performed colony-forming units counting and analysis. SK and TK performed flow cytometry analysis and FluoroSpot. SK, TK, and TW performed data and statistical analysis. SK, JTG, PN, TK, JRC, DQ, KC, AS, YH, SKA, RT, SC, CD, TW, CS, RBM and PCK interpreted the data and edited the manuscript.

## Notes

**Conflict of Interest:** The authors have declared that no conflict of interest exists.

### Competing Interest Statement

The authors have declared no competing interest.

