## Supplementary Figures and Tables for "An intranasal stringent response vaccine targeting dendritic cells as a novel adjunctive therapy against tuberculosis"

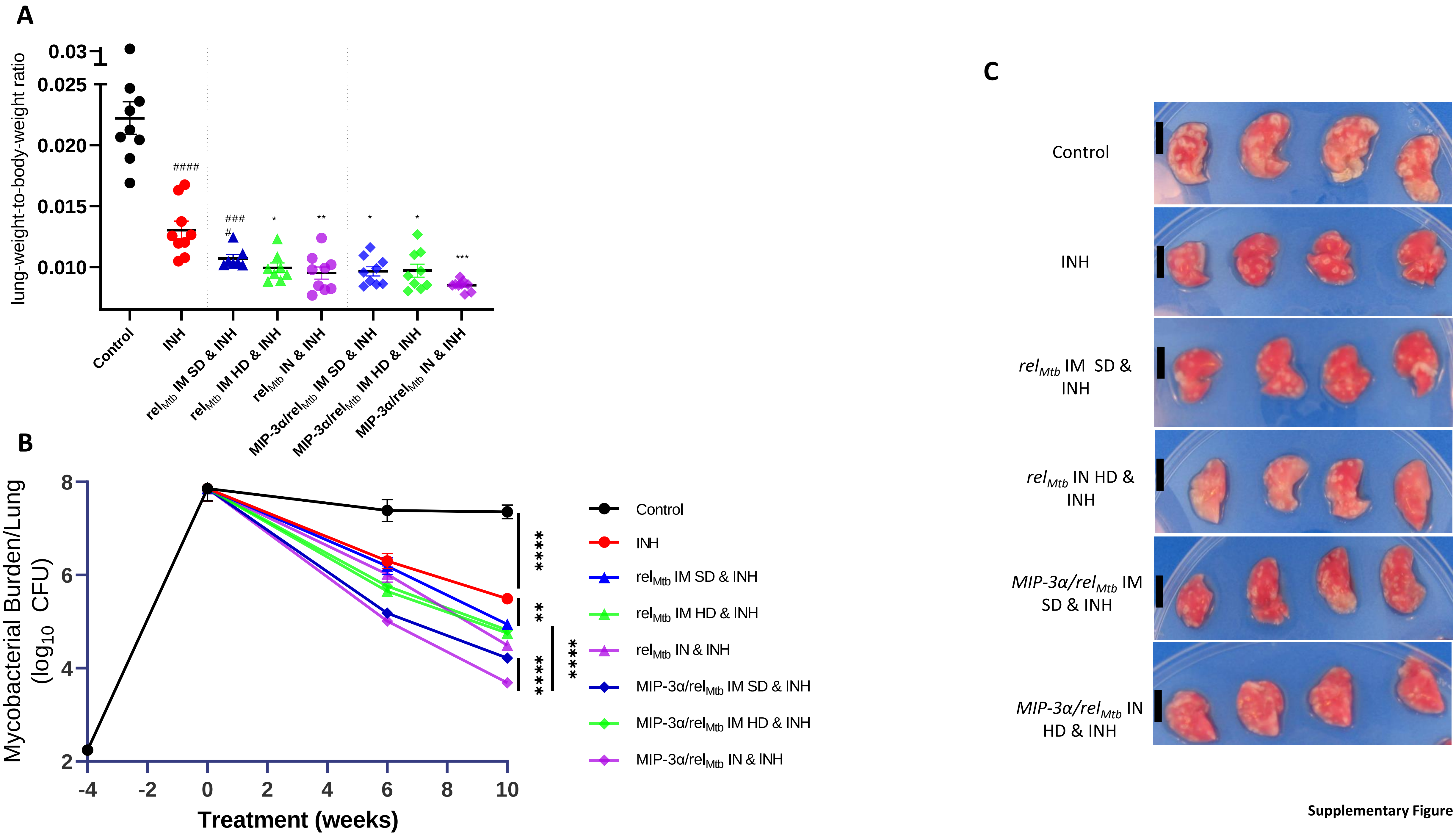


Supplementary Figure 1

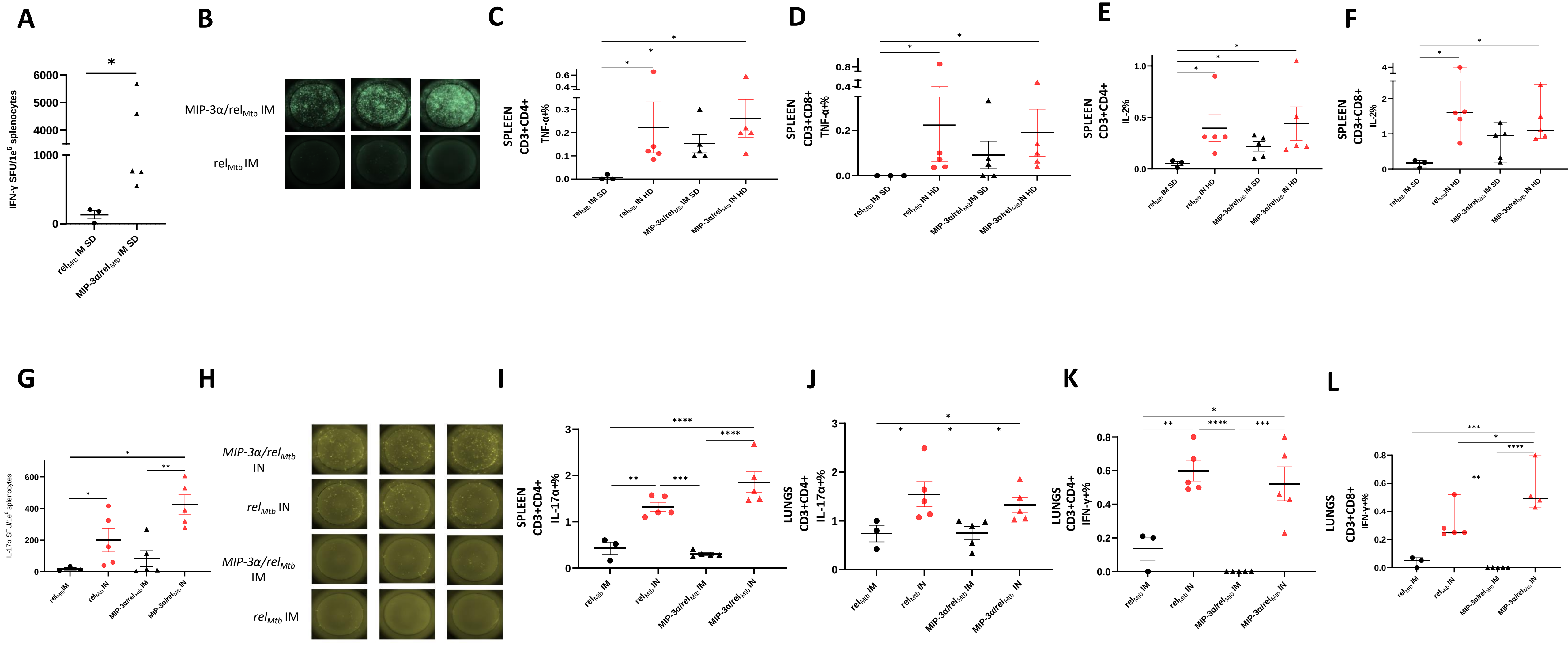


Supplementary Figure 2

#### Supplementary Figure Legends

**Supplementary Figure 1.** (A) Normalized mean mouse lung weights at 10 weeks (B) Lung mycobacterial burden at: implantation (-4 weeks); initiation of treatment (0 weeks); and at 6 weeks and 10 weeks after the initiation of treatment. (C) Gross pathology of representative lungs per experimental group; black line represents 1 cm, TB: tuberculosis, IM: Intramuscular, IN: Intranasal, SD: Standard Dose, HD: High Dose, CFU: colony-forming units. *** = significant difference from control and INH (at least P< 0.001), ** significant difference from control and INH (at least P<0.01), * significant difference from control and INH (at least P<0.05), #### significant difference from control only (P<0.0001).

**Supplementary Figure 2: T-cell responses in non-infected murine tissues after vaccination (immunogenicity, non-challenged, animal study).** (A) $Rel_{Mtb}$-specific IFN-γ response in spleens between IM vaccination with *MIP-3α/rel*$_{Mtb}$ *vs rel*$_{Mtb}$ , as assessed by FluoroSpot . (B) Representative pictures of FluoroSpot are shown per group. Each triplicate represents three different mice. (C) and (D) $Rel_{Mtb}$-specific TNF-α producing CD4+ and CD8+ T cells in cells derived from spleens among the different vaccination groups as assessed by flow cytometry. (E) and (F) $Rel_{Mtb}$-specific IL-2 producing CD4+ and CD8+ T cells among in cells derived from spleens among the different vaccination groups as assessed by flow cytometry. (G) $Rel_{Mtb}$-specific IL-17α response in spleens among different vaccination groups as assessed by FluoroSpot. (H) Representative pictures of FluoroSpot are shown per group. Each triplicate represents three different mice. (I) $Rel_{Mtb}$-specific IL-17α producing CD4+ T cells in cells derived from spleens among the different vaccination groups as assessed by flow cytometry (J) $Rel_{Mtb}$-specific IL-17α producing CD4+ T cells in cells derived from lungs among the different vaccination groups as assessed by flow cytometry (K) and (L) $Rel_{Mtb}$-specific TNF-α producing CD4+ and CD8+ T cells in cells derived from spleens among the different vaccination groups as assessed by flow cytometry. (E) and (F) $Rel_{Mtb}$-specific IFN-γ producing CD4+ and CD8+ T cells in cells derived from lungs among the different vaccination groups as assessed by flow cytometry. SFU: Spot Forming Unit, IM: Intramuscular, IN: Intranasal. Y-axis scales are different among cytokines and between tissues in order to better demonstrate differences between groups where cytokine expression levels were lower.

**Supplementary Table 1.** Lung mycobacterial burden of *Mtb*-infected mice at 10 weeks post infection. INH: Isoniazid, IM: Intramuscular, IN: Intranasal, SD: Standard dose, HD: High dose

| LUNG MYCOBACTERIAL BURDEN AT 10 WEEKS | |
|---|---|
| **Comparisons** | **Adjusted P Value** |
| Control vs. INH | <0.0001 |
| Control vs. $rel_{Mtb}$ IM SD & INH | <0.0001 |
| Control vs. $rel_{Mtb}$ IM HD & INH | <0.0001 |
| Control vs. $rel_{Mtb}$ IN HD & INH | <0.0001 |
| Control vs. MIP-3α/$rel_{Mtb}$ IM SD & INH | <0.0001 |
| Control vs. MIP-3α/$rel_{Mtb}$ IM HD & INH | <0.0001 |
| Control vs. MIP-3α/$rel_{Mtb}$ IN HD & INH | <0.0001 |
| INH vs. $rel_{Mtb}$ IM SD & INH | 0.0076 |
| INH vs. $rel_{Mtb}$ IM HD & INH | <0.0001 |
| INH vs. $rel_{Mtb}$ IN HD & INH | <0.0001 |
| INH vs. MIP-3α/$rel_{Mtb}$ IM SD & INH | <0.0001 |
| INH vs. MIP-3α/$rel_{Mtb}$ IM HD & INH | 0.0002 |
| INH vs. MIP-3α/$rel_{Mtb}$ IN HD & INH | <0.0001 |
| $rel_{Mtb}$ IM SD & INH vs. $rel_{Mtb}$ IM HD & INH | 0.8952 |
| $rel_{Mtb}$ IM SD & INH vs. $rel_{Mtb}$ IN HD & INH | 0.0052 |
| $rel_{Mtb}$ IM SD & INH vs. MIP-3α/$rel_{Mtb}$ IM SD & INH | 0.0001 |
| $rel_{Mtb}$ IM SD & INH vs. MIP-3α/$rel_{Mtb}$ IM HD & INH | 0.9901 |
| $rel_{Mtb}$ IM SD & INH vs. MIP-3α/$rel_{Mtb}$ IN HD & INH | <0.0001 |
| $rel_{Mtb}$ IM HD & INH vs. $rel_{Mtb}$ IN HD & INH | 0.5415 |
| $rel_{Mtb}$ IM HD & INH vs. MIP-3α/$rel_{Mtb}$ IM SD & INH | 0.0050 |
| $rel_{Mtb}$ IM HD & INH vs. MIP-3α/$rel_{Mtb}$ IM HD & INH | 0.9997 |
| $rel_{Mtb}$ IM HD & INH vs. MIP-3α/$rel_{Mtb}$ IN HD & INH | <0.0001 |

| | |
|---|---|
| $rel_{Mtb}$ IN HD & INH vs. MIP-3α/$rel_{Mtb}$ IM SD & INH | 0.4858 |
| $rel_{Mtb}$ IN HD & INH vs. MIP-3α/$rel_{Mtb}$ IM HD & INH | 0.2519 |
| $rel_{Mtb}$ IN HD & INH vs. MIP-3α/$rel_{Mtb}$ IN HD & INH | <0.0001 |
| MIP-3α/$rel_{Mtb}$ IM SD & INH vs. MIP-3α/$rel_{Mtb}$ IM HD & INH | 0.0010 |
| MIP-3α/$rel_{Mtb}$ IM SD & INH vs. MIP-3α/$rel_{Mtb}$ IN HD & INH | 0.0058 |
| MIP-3α/$rel_{Mtb}$ IM HD & INH vs. MIP-3α/$rel_{Mtb}$ IN HD & INH | <0.0001 |

**Supplementary Table 2. T-cell responses in *Mtb*-infected murine tissues 6 weeks post treatment initiation.** INH: Isoniazid, IM: Intramuscular, IN: Intranasal.

| T-cell responses in *Mtb*-infected murine tissues | | | | | | | | | | |
|---|---|---|---|---|---|---|---|---|---|---|
| | SPLEEN | | | | | | | LUNG | | |
| | IFN-γ | | TNF-α | | IL-2 | | IL-17α | IFN-γ | | IL-17α |
| | CD4+ T | CD8+T | CD4+ T | CD8+T | CD4+ T | CD8+T | CD4+ T | CD4+ T | CD8+T | CD4+ T |
| **Tukey's Multiple Comparisons Test** | **Adjusted P Value** | | | | | | | | | |
| Control vs. INH | 0.3348 | 0.2939 | >0.9999 | >0.9999 | >0.9999 | >0.9999 | >0.9999 | 0.9999 | >0.9999 | 0.9990 |
| Control vs. $rel_{Mtb}$ IM & INH | 0.3741 | 0.0838 | 0.9251 | 0.9815 | 0.9871 | 0.9949 | 0.7119 | >0.9999 | >0.9999 | 0.9616 |
| Control vs. $rel_{Mtb}$ IN & INH | 0.0557 | <0.0001 | 0.0010 | 0.0972 | 0.0002 | <0.0001 | 0.0025 | 0.0447 | 0.0057 | <0.0001 |
| Control vs. MIP-3α/$rel_{Mtb}$ IM & INH | <0.0001 | <0.0001 | 0.0002 | 0.2972 | 0.0007 | <0.0001 | 0.4208 | 0.6696 | 0.4597 | 0.7090 |
| Control vs. MIP-3α/$rel_{Mtb}$ IN & INH | <0.0001 | <0.0001 | <0.0001 | <0.0001 | <0.0001 | <0.0001 | <0.0001 | <0.0001 | <0.0001 | <0.0001 |
| INH vs. $rel_{Mtb}$ IM & INH | >0.9999 | 0.9876 | 0.9751 | 0.9894 | 0.9856 | 0.9941 | 0.7898 | 0.9998 | >0.9999 | 0.9975 |
| INH vs. $rel_{Mtb}$ IN & INH | 0.9465 | 0.0020 | 0.0020 | 0.1159 | 0.0002 | <0.0001 | 0.0037 | 0.0805 | 0.0105 | <0.0001 |
| INH vs. MIP-3α/$rel_{Mtb}$ IM & INH | <0.0001 | <0.0001 | 0.0002 | 0.3386 | 0.0007 | <0.0001 | 0.5046 | 0.8121 | 0.5964 | 0.8957 |
| INH vs. MIP-3α/$rel_{Mtb}$ IN & INH | <0.0001 | <0.0001 | <0.0001 | <0.0001 | <0.0001 | <0.0001 | <0.0001 | <0.0001 | <0.0001 | <0.0001 |
| $rel_{Mtb}$ IM & INH vs. $rel_{Mtb}$ IN & INH | 0.9268 | 0.0123 | 0.0167 | 0.3597 | 0.0018 | <0.0001 | 0.1052 | 0.0439 | 0.0063 | <0.0001 |
| $rel_{Mtb}$ IM & INH vs . MIP-3α/$rel_{Mtb}$ IM & INH | <0.0001 | <0.0001 | 0.0076 | 0.7128 | 0.0050 | <0.0001 | 0.9970 | 0.6648 | 0.4816 | 0.9900 |
| $rel_{Mtb}$ IM & INH vs. MIP-3α/$rel_{Mtb}$ IN & INH | <0.0001 | <0.0001 | <0.0001 | <0.0001 | <0.0001 | <0.0001 | <0.0001 | <0.0001 | <0.0001 | <0.0001 |
| $rel_{Mtb}$ IN & INH vs. MIP-3α/$rel_{Mtb}$ IM & INH | <0.0001 | <0.0001 | 0.5341 | 0.9921 | 0.9993 | 0.9451 | 0.2627 | 0.6451 | 0.3761 | <0.0001 |
| $rel_{Mtb}$ IN & INH vs. MIP-3α/$rel_{Mtb}$ IN & INH | <0.0001 | <0.0001 | 0.0003 | <0.0001 | 0.5818 | 0.8022 | <0.0001 | 0.0340 | 0.0064 | 0.1005 |
| MIP-3α/$rel_{Mtb}$ IM & INH vs. MIP-3α/$rel_{Mtb}$ IN & INH | 0.717 | 0.9929 | <0.0001 | <0.0001 | 0.3635 | 0.2645 | <0.0001 | 0.0003 | <0.0001 | <0.0001 |

**Supplementary Table 3. T-cell responses in uninfected murine tissues 6 weeks after primary vaccination.** IM: Intramuscular, IN: Intranasal. PBMC: peripheral blood mononuclear cells.

| T-cell responses in uninfected murine tissues | | | | |
|---|---|---|---|---|
| | **Draining Lymph Nodes** | **PBMCs** | **PBMCs** | |
| | IL-17α | IL-17α | TNF-α | |
| | CD4+ T | CD4+ T | CD4+ T | |
| **Tukey's Multiple Comparisons Test** | | | Day 28 | Day 42 |
| | **Adjusted P Value** | | | |
| rel$_{Mtb}$ IM vs. rel$_{Mtb}$ IN | 0.0003 | 0.0007 | 0.085 | 0.036 |
| rel$_{Mtb}$ IM vs. MIP-3α/rel$_{Mtb}$ IM | 0.0357 | 0.0357 | 0.036 | 0.036 |
| rel$_{Mtb}$ IM vs. MIP-3α/rel$_{Mtb}$ IN | 0.0003 | 0.0265 | 0.025 | 0.010 |
| rel$_{Mtb}$ IN vs. MIP-3α/rel$_{Mtb}$ IM | 0.0033 | 0.0023 | 0.9889 | 0.9339 |
| rel$_{Mtb}$ IN vs. MIP-3α/rel$_{Mtb}$ IN | >0.9999 | 0.1522 | 0.5171 | 0.9462 |
| MIP-3α/rel$_{Mtb}$ IM vs. MIP-3α/rel$_{Mtb}$ IN | 0.0034 | 0.0079 | 0.123 | 0.036 |

### Supplementary Appendix

#### DNA SEQUENCE

#### Mouse Codon Optimized MIP-3α/Rel$_{Mtb}$

##### Top Strand Bases

GCCGCTAGCAACTTCGACTGCTGTCTGGGATACACAGATAGAATCCTGCACCCAAAGTTCATCGTGGGCTTTACCAGACAGCT
GGCCAACGAGGGATGCGACATCAACGCTATCATCTTTCACACCAAGAAGAAGCTGAGCGTGTGCGCCAACCCCAAGCAGACA
TGGGTGAAGTACATCGTGCGGCTGCTGAGCAAGAAGGTGAAGAACATG **(MIP-3α)**

GGACCAGGACCTGGACCAGGACCAGGACCTCAGGCGCCGAAGAGTCTCGAGgctagc **(Linker)**

ACCGCCCAGAGGTCTACCACAAACCCTGTGCTGGAGCCACTGGTGGCAGTCCACAGGGAGATCTACCCCAAGGCCGATCTGA
GCATCCTGCAGAGGGCATATGAGGTGGCAGACCAGAGGCACGCCAGCCAGCTGCGCCAGTCCGGCGATCCTTACATCACACA
CCCACTGGCCGTGGCCAATATCCTGGCCGAGCTGGGCCTGGACACCACAACCCTGGTGGCCGCCCTGCTGCACGACACCGTGG
AGGATACAGGCTATACCCTGGAGGCCCTGACAGAGGAGTTCGGAGAGGAAGTGGGACACCTGGTGGACGGAGTGACCAAGC
TGGATAGGGTGGTGCTGGGCTCCGCCGCAGAGGGAGAGACAATCAGAAAGATGATCACAGCAATGGCCAGGGACCCCAGGG
TGCTGGTCATCAAGGTGGCCGACCGGCTGCACAACATGAGGACCATGAGATTCCTGCCACCTGAGAAGCAGGCAAGGAAGGC
CAGGGAGACACTGGAAGTGATCGCACCACTGGCCCACAGGCTGGGAATGGCCTCTGTGAAGTGGGAGCTGGAGGACCTGAG
CTTTGCCATCCTGCACCCTAAGAAGTACGAGGAGATCGTGCGGCTGGTGGCAGGAAGGGCACCAAGCAGAGATACCTATCTG
GCCAAGGTGCGCGCCGAGATCGTGAATACACTGACCGCCTCTAAGATCAAGGCCACAGTGGAGGGCAGGCCCAAGCACTACT
GGAGCATCTATCAGAAGATGATCGTGAAGGGCAGAGACTTCGACGATATCCACGATCTGGTGGGCGTGAGAATCCTGTGCGA
CGAGATCCGCGATTGTTACGCAGCAGTGGGAGTGGTGCACAGCCTGTGGCAGCCAATGGCAGGCCGGTTTAAGGACTATATC
GCCCAGCCCCGCTACGGCGTGTATCAGTCCCTGCACACAACCGTGGTGGGACCAGAGGGCAAGCCTCTGGAGGTGCAGATCC
GGACCCGCGATATGCACAGGACAGCAGAGTACGGAATCGCAGCACACTGGAGGTATAAGGAGGCCAAGGGCAGAAACGGC
GTGCTGCACCCTCACGCAGCAGCAGAGATCGACGATATGGCCTGGATGAGGCAGCTGCTGGACTGGCAGAGGGAGGCAGCC
GATCCCGGAGAGTTCCTGGAGTCTCTGCGCTACGACCTGGCCGTGCAGGAGATCTTCGTGTTTACCCCTAAGGGCGACGTGAT
CACACTGCCCACCGGCAGCACACCTGTGGATTTTGCCTATGCAGTGCACACAGAAGTGGGACACAGGTGCATCGGAGCCCGG
GTGAACGGCCGCCTGGTGGCCCTGGAGCGCAAGCTGGAGAATGGCGAGGTGGTGGAGGTGTTTACCAGCAAGGCACCAAAC
GCAGGACCCTCCAGAGACTGGCAGCAGTTCGTGGTGTCCCCAAGGGCCAAGACCAAGATCAGACAGTGGTTTGCCAAGGAGA
GGAGAGAGGAGGCCCTGGAGACAGGCAAGGATGCCATGGCCCGGGAGGTGCGGAGGGGAGGCCTGCCCCTGCAGCGCCTG
GTGAATGGAGAGTCTATGGCAGCAGTGGCCAGGGAGCTGCACTACGCAGACGTGAGCGCCCTGTATACCGCAATCGGAGAG
GGACACGTGTCCGCCAAGCACGTGGTGCAGAGACTGCTGGCCGAGCTGGGAGGAATCGATCAGGCCGAGGAGGAGCTGGCC
GAGAGGTCTACCCCAGCCACAATGCCCAGGAGGCCCAGATCTACCGACGATGTGGGCGTGAGCGTGCCAGGAGCACCAGGC
GTGCTGACCAAGCTGGCCAAGTGCTGTACACCAGTGCCCGGCGACGTGATCATGGGATTCGTGACAAGGGGCGGAGGCGTGT
CCGTGCACAGAACCGATTGTACAAACGCAGCCTCTCTGCAGCAGCAGGCAGAGAGGATCATCGAGGTGCTGTGGGCCCCTTC
CCCAAGCTCCGTGTTTCTGGTGGCCATCCAGGTGGAGGCCCTGGACAGGCACAGACTGCTGTCTGATGTGACCAGAGCCCTGG
CCGACGAGAAAGTGAATATCCTGTCTGCCAGCGTGACAACCTCCGGCGACAGGGTGGCCATCAGCAGGTTCACCTTCGAGAT
GGGCGATCCTAAGCACCTGGGCCACCTGCTGAACGCCGTGAGGAATGTGGAGGGCGTGTACGACGTGTATAGAGTGACCTCC
GCCGCC **(Mouse codon optimized Rel$_{Mtb}$)**

### DNA SEQUENCE

#### Rel$_{Mtb}$ (non-codon optimized)

##### Top Strand Bases

GTGGCCGAGGACCAGCTCACGGCGCAAGCGGTTGCACCGCCCACGGAGGCTTCTGCGGCTCTCGA
GCCCGCTCTCGAGACGCCCGAGTCGCCGGTCGAGACTCTTAAGACCAGCATCAGCGCGTCGCGTC
GGGTGCGGGCCCGATTGGCCCGGCGGATGACCGCCCAGCGCAGCACCACCAATCCGGTGCTCGA
GCCGTTGGTGGCGGTGCACCGGGAGATCTATCCCAAGGCCGACCTGTCGATCTTGCAGCGAGCCT
ACGAGGTCGCTGACCAAAGGCATGCCAGCCAGTTGCGGCAGTCCGGTGATCCCTACATCACCCACC
CGTTGGCCGTTGCCAACATTCTGGCCGAGTTGGGCATGGACACCACCACTTTGGTGGCCGCGCTGC
TGCACGACACCGTCGAGGACACCGGTTACACCCTGGAGGCGTTGACCGAGGAATTCGGCGAAGAG
GTGGGCCATCTCGTCGACGGGGTGACCAAGCTGGATCGGGTGGTGTTGGGCAGCGCCGCCGAAG
GCGAGACTATTCGCAAGATGATCACCGCGATGGCCCGCGATCCGCGGGTGCTGGTGATAAAGGTG
GCTGACCGGTTACACAACATGCGCACCATGCGCTTCTTGCCGCCGGAGAAGCAGGCCCGCAAGGC
CCGTGAGACGTTGGAAGTCATTGCACCCCTGGCGCATCGGCTGGGCATGGCCAGCGTCAAGTGGG
AGTTGGAGGACCTGTCCTTCGCGATCCTGCATCCCAAGAAGTACGAGGAGATCGTCCGGCTGGTCG
CCGGTCGCGCGCCGTCCCGGGACACCTACCTGGCCAAGGTGCGTGCCGAAATCGTCAACACGCTG
ACCGCGTCGAAGATCAAGGCGACGGTGGAGGGCCGCCCCAAGCACTATTGGTCGATCTACCAGAA
GATGATCGTTAAGGGCCGCGACTTCGACGACATCCACGACCTGGTCGGTGTGCGCATCCTGTGCGA
CGAAATCCGGGACTGCTACGCGGCTGTCGGCGTAGTGCATTCGCTATGGCAGCCGATGGCGGGTC
GGTTCAAGGACTACATCGCCCAGCCCAGATACGGTGTGTACCAGTCACTGCACACCACTGTGGTCG
GGCCTGAGGGCAAGCCGCTGGAAGTGCAGATCCGTACCCGCGACATGCACCGCACCGCCGAATAC
GGCATCGCCGCGCATTGGCGCTACAAAGAAGCCAAGGGCCGCAACGGTGTTCTTCATCCGCATGCC
GCCGCGGAGATCGACGACATGGCCTGGATGCGTCAGCTGCTCGACTGGCAACGTGAGGCGGCCGA
CCCCGGTGAGTTCTTGGAATCATTGCGCTACGACCTTGCGGTGCAAGAGATTTTCGTGTTTACCCCC
AAGGGCGACGTGATCACGCTGCCAACCGGTTCGACGCCGGTGGACTTCGCTTACGCGGTGCACAC
AGAGGTGGGCCACCGCTGCATCGGCGCCCGAGTGAACGGCCGGTTGGTAGCGCTGGAACGCAAG
CTGGAAAACGGAGAAGTTGTCGAGGTTTTCACGTCCAAGGCGCCGAACGCCGGGCCGTCGCGGGA
CTGGCAGCAGTTCGTGGTGTCGCCGCGCGCAAAGACGAAGATCCGCCAGTGGTTCGCCAAGGAGC
GGCGTGAGGAGGCGTTGGAGACCGGTAAGGATGCGATGGCCCGCGAGGTGCGCCGCGGTGGACT
TCCGTTGCAGCGCTTGGTCAATGGTGAGTCCATGGCGGCGGTGGCCCGCGAGCTGCACTACGCGG
ACGTGTCAGCACTCTATACCGCCATCGGTGAGGGGCACGTGTCGGCGAAACACGTCGTGCAGCGG
TTGTTGGCCGAGCTCGGCGGTATCGACCAGGCGGAAGAGGAACTCGCCGAGCGGTCCACGCCGGC
GACCATGCCGCGGCGCCCACGCAGCACCGACGATGTCGGGGTCTCCGTCCCCGGCGCCCCGGGC
GTGCTGACCAAGCTGGCCAAGTGCTGCACGCCGGTTCCGGGCGATGTGATTATGGGGTTCGTCACC
CGTGGCGGCGGGGTCAGTGTGCACCGCACCGACTGCACCAACGCCGCATCGCTGCAGCAGCAGG
CCGAGCGCATCATCGAGGTGCTATGGGCGCCGTCGCCGTCGTCGGTGTTTCTGGTGGCAATCCAG
GTCGAGGCACTCGACCGGCACCGGCTGCTGTCGGATGTGACGCGCGCACTGGCCGACGAGAAGGT
CAATATCCTGTCCGCGTCGGTCACCACTTCGGGGGACCGGGTGGCGATCAGTCGATTCACCTTCGA
GATGGGTGACCCCAAGCACCTCGGGCACCTGCTCAACGCCGTCCGCAACGTCGAAGGTGTCTACG
ACGTCTACCGGGTGACCTCGGCCGCG
